# Coevolutionary cycling in allele frequencies and the evolution of virulence

**DOI:** 10.1101/2025.05.22.655598

**Authors:** Yoon Soo Kim, Ben Ashby

**Affiliations:** Department of Mathematics, Simon Fraser University, Burnaby, BC, Canada

**Author notes:** **Data availability** Source code is available in the *Supplementary Material* and at https://github.com/dxanielkm/Ashby_and_Kim_2025/tree/main. **Author contributions** Investigation, software, visualization: YSK; Conceptualization, supervision, methodology: BA; Writing: YSK and BA.

## Abstract

Coevolutionary cycling in allele frequencies due to negative frequency-dependent selection—sometimes referred to as Red Queen Dynamics—is a key potential outcome of host-parasite coevolution. While many theoretical studies have focused on understanding the consequences of coevolutionary cycling for the evolution of sex and recombination, little is known about the impact of coevolutionary cycling on the evolution of other life history traits. It is therefore currently unknown how coevolutionary cycling in allele frequencies affects the evolution of key disease characteristics, such as virulence. Here, we combine population genetic and quantitative genetic approaches to theoretically determine the impacts of coevolutionary cycling in allele frequencies on the evolution of virulence in a free-living parasite. By varying the level of genetic specificity required for infection while controlling for the average infection rate, we induce coevolutionary cycles and examine their effects on virulence evolution. We show that coevolutionary cycling does indeed have a strong impact on virulence evolution, with more specific infection genetics and higher allelic diversity generally driving larger and more rapid cycles in allele frequencies, leading to selection for higher virulence. Our research provides new fundamental insights into the relationship between coevolutionary cycling and the evolution of virulence.

## 1 Introduction

Host-parasite interactions are ubiquitous in nature, with antagonistic coevolution between species playing a critical role in shaping biodiversity and the evolution of a wide range of traits. Cycling in allele frequencies due to negative frequency-dependent selection (often used synonymously with Red Queen Dynamics) is a hallmark, but notably, not the only outcome, of host-parasite coevolution (Sasaki 2000; Best et al. 2010; Brockhurst et al. 2014; Ashby 2020). If parasites tend to specialise on certain hosts, they will typically adapt to infect the most common genotypes, leading to a potential advantage for rare host alleles. Rare host alleles are then expected to increase in frequency, with corresponding parasite alleles following suit, tracking the most common host alleles. Coevolutionary cycling can potentially persist indefinitely, maintaining genetic diversity through time, and has been observed or inferred in a variety of taxa, including animal hosts and bacterial pathogens (Decaestecker et al. 2007; Papkou et al. 2019) or trematode parasites (Dybdahl and Lively 1998), bacterial hosts and viral pathogens (Gómez and Buckling 2011; Hall et al. 2011), and plants and fungal pathogens (Chaboudez and Burdon 1995). These dynamics are thought to be important for a wide range of biological phenomena, from the evolution of sex and recombination (Hamilton 1980; Lively 2010), to parasite-mediated sexual selection (Hamilton and Zuk 1982), to local adaptation (Koskella 2014; Ashby and Boots 2017).

The theoretical literature on host-parasite coevolution (reviewed in Ashby et al. (2019) and Buckingham and Ashby (2022)) largely focuses on either population genetics (e.g., cycling in allele frequencies (Frank 1993; Sasaki 2000; Agrawal and Lively 2002; Ashby and Boots 2017)) or the coevolution of quantitative or life-history traits (e.g., host resistance and parasite virulence (Sasaki and Godfray 1999; Nuismer et al. 2005; Best et al. 2010; Ashby and Boots 2015)). While there is a rich body of theory in both areas, there is limited overlap between them. In particular, very little is known about how coevolution among genes that determine whether and to what extent a parasite can infect a given host (infection genetics), affect the evolution of key disease-related traits, such as virulence. As far as we are aware, only two theoretical studies have explored this question previously (Kirchner and Roy 2002; Gandon et al. 2002). Kirchner and Roy (2002) explored a two-host, two-pathogen model with variable specificity to determine its impact on the evolution of pathogen infectiousness and virulence. They showed that at sufficiently high levels of specificity, less lethal and more lethal parasite strains are able to co-exist due to negative frequency-dependent selection. However, their model only considered the two-host, two-pathogen case with fixed levels of virulence, rather than allowing virulence to evolve freely. Gandon et al. (2002) explored how the Gene-for-Gene (GFG) framework of infection genetics affects the evolution of parasite virulence. The GFG framework, inspired by flax rust (Flor 1956), assumes that hosts vary in the range of parasites that they can resist, and parasites vary in the range of hosts that they can infect. In the simplest case, one host genotype is susceptible to both parasites but the other host genotype is resistant to a single parasite. Resistance and broader host range are often associated with pleiotropic fitness costs, which can maintain polymorphism at a stable equilibrium or lead to coevolutionary cycling (Buckingham and Ashby 2022). Gandon et al. (2002) found that GFG coevolution decreases the evolutionarily stable level of virulence compared to the absence of coevolution, but only when superinfection occurs (i.e., when a new infection can replace an old infection). However, the authors focused on one type of genetic framework (the GFG model with two host and two parasite genotypes) and scenarios where the populations are at a stable equilibrium, rather than exhibiting coevolutionary cycling.

Here, we use eco-evolutionary simulations of a generalisation of the ‘Matching Alleles’ (MA) framework of infection genetics (Frank 1993; Grosberg and Hart 2000) to explore how coevolutionary cycling impacts the evolution of parasite virulence. By varying the strength of specificity between host and parasite genotypes and the diversity of susceptibility and infectivity alleles while controlling for the mean infection rate as specificity and genetic diversity vary, we show that more rapid and higher amplitude coevolutionary cycles select for higher parasite virulence.

## 2 Methods

### 2.1 Model description

We consider the coevolutionary dynamics of a well-mixed host population and free-living parasite population. Free-living parasites such as bacteriophages, protozoa, and helminths are common in nature, but are relatively understudied in theoretical evolutionary biology, with most models focusing on non-free-living parasites. Furthermore, free-living parasites are known to produce sustained population cycles under certain conditions, which allows us to focus on long-term evolutionarily stable levels of virulence under persistent coevolutionary cycling. Both host and parasite populations are asexual and haploid with host susceptibility to infection determined by interactions at a single genetic locus, with *n* alleles in each population. We refer to these alleles as ‘susceptibility’ alleles for the host and ‘infectivity’ alleles for the parasite. Note that what we refer to as infectivity alleles are often referred to as ‘virulence’ alleles in the literature, especially in the context of plant-pathogens, but here we make the important distinction between the ability to infect a host (infectivity) and the severity of disease caused to the host (virulence). Parasites also have a second locus, with *m* alleles, which determines their level of virulence in the form of the disease-associated mortality rate. There are therefore *n* host genotypes and *mn* possible parasite genotypes. Let *S*_*i*_ be the density of susceptible hosts with susceptibility allele *i, I*_*ijk*_ be the density of hosts with susceptibility allele *i* infected by parasites with infectivity allele *j* and virulence allele *k*, and *P*_*jk*_ be the density of free-living parasites with infectivity allele *j* and virulence allele *k*. We assume that coinfection and superinfection do not occur.

Hosts with susceptibility allele *i* reproduce at rate *bS*_*i*_(1 *− qN*), where *b* represents the intrinsic birth rate, *q* is the strength of density-dependent competition, and 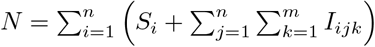 is the total host population density. All hosts have a natural mortality rate of *d*, and infected hosts have an additional disease-associated mortality rate *α*_*k*_ (which depends on the virulence allele of the parasite, *k*) and recovery rate *γ*. Infected hosts shed parasites at a constant rate *θ*_*k*_ and free-living parasites die or are removed from the environment at a constant rate of *δ*. We assume that parasites experience a trade-off between their shedding rate and virulence such that 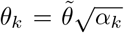, where 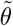 scales the shedding rate. This assumes that a higher shedding rate is associated with harvesting more host resources (e.g., infecting more cells), leading to higher virulence. However, there are diminishing returns for parasites with higher shedding rates due to greater damage to the host which results in an accelerating host mortality rate. While many studies on the evolution of virulence assume a transmission-virulence trade-off, a shedding rate-virulence trade-off is more mechanistically justified in a model with free-living parasites as the production of free-living parasite propagules is directly linked to the harm caused to the host (e.g., Lion and Gandon (2022)).

Free-living parasites with infectivity allele *j* and virulence allele *k* infect uninfected hosts with susceptibility allele *i* at rate *β*(*n*)*Q*_*ij*_(*n, s*)*P*_*jk*_*S*_*i*_, where *β*(*n*) is the baseline infection rate and *Q*_*ij*_(*n, s*) is the *n × n* ‘specificity matrix’ with specificity parameter *s ∈* [0, 1] (similar to Kirchner and Roy (2002)). The specificity matrix determines the extent to which parasites specialise on hosts with matching alleles, with:

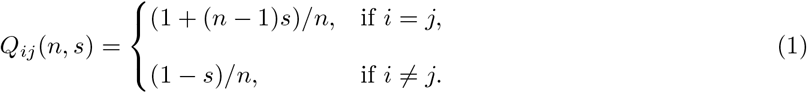

We construct this matrix so that as the specificity parameter *s* increases, the diagonal elements (representing matching alleles between hosts and parasites) increase, while the off-diagonal elements decrease (representing mismatching alleles). When *s* = 0, there is no specificity and so all elements of *Q*_*ij*_(*n*, 0) are equal to 1*/n* (i.e., all hosts are equally susceptible to all parasites). When *s* = 1, the specificity matrix is equal to the identity matrix (i.e., hosts are only susceptible to parasites with matching alleles). Note that the rows and columns of the specificity matrix always sum to one, which ensures that specificity has no effect on the average infection rate. Furthermore, as there are no differences in life-history traits between host genotypes, the mean frequency of each host susceptibility allele is 1*/n* when all parasites have the same virulence. We therefore set the baseline infection rate, *β*(*n*) = *β*_0_*n*, to be the product of an underlying infection rate, *β*_0_, and the number of possible infectivity alleles, *n*, which ensures that mean disease prevalence does not decrease as the number of infectivity alleles increases. Hence, variation in specificity (*s*) or the number of susceptibility and infectivity alleles (*n*) does not affect the average infection rate when the populations are at equilibrium, and so any impact on the evolution of virulence is entirely due to coevolutionary cycling of susceptibility and infectivity alleles.

The eco-evolutionary dynamics are given by the following set of ordinary differential equations (ODEs):

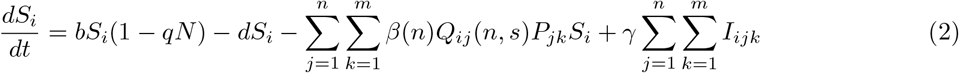

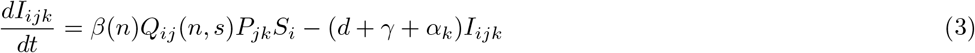

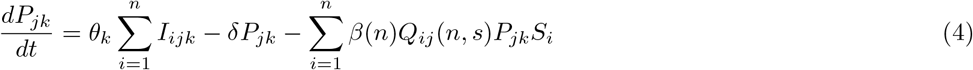

Without loss of generality, we set *d* = 1 and *q* = 1 throughout our analysis as the model can be readily non-dimensionalised to remove these parameters.

### 2.2 Analysis

We use hybrid simulations with deterministic and stochastic components to analyze the impacts of coevolutionary cycles in allele frequencies on the evolution of parasite virulence (the full simulation algorithm and source code are available in the *Supplementary Materials* and at https://github.com/dxanielkm/Ashby_and_Kim_2025/tree/main). Each simulation is initialised with all susceptibility and infectivity alleles present at arbitrary population densities, and a single virulence phenotype for all parasites.

The simulations are separated into ecological and evolutionary timescales. Within each evolutionary time step, the ecological dynamics of extant hosts and parasites are described by equations (2)-(4). The ecological dynamics are progressed deterministically for a fixed number of time units, *T*_*eco*_. The duration of ecological dynamics effectively controls the mutation rate for the virulence locus, with smaller values of *T*_*eco*_ corresponding to a higher mutation rate and larger values to a lower mutation rate. The ecological dynamics are then paused and any free-living parasite compartment (*P*_*jk*_) that is below a density of *ϵ* = 10^*−*9^ is set to 0. We then introduce a rare mutant parasite with a slightly different virulence phenotype, *α*_*mut*_, to one of the resident phenotypes, *α*_*res*_. The progenitor for the mutant is chosen probabilistically using a cumulative distribution function of the free-living parasite classes (see *Supplementary Materials* for the full mutation algorithm). The period of coevolutionary cycles is generally not a multiple of *T*_*eco*_ and also varies as virulence evolves, which removes any bias in the timing of mutations with the current state of the population (also confirmed in preliminary simulations). The ecological dynamics with the new mutant are then progressed again by *T*_*eco*_ time units and the process repeats for a total of *T*_*evo*_ = 4, 000 evolutionary time steps (this duration was sufficient for convergence to an evolutionarily stable level of virulence in all simulations).

The duration for the ecological dynamics and the extinction thresholds are fairly arbitrary, with preliminary simulations confirming that a wide range of values produce similar results, provided *T*_*eco*_ is not too small (i.e., the mutation rate is not too high) and *ϵ* is not too large (i.e., only rare phenotypes are removed from the system). We therefore fix *T*_*eco*_ = 400 throughout our analysis, but show in the *Supplementary Materials* that faster or slower mutation rates (i.e., smaller or larger values of *T*_*eco*_) do not qualitatively affect our results (Figs. S1, S4-S5). Note that the extinction threshold is only included to improve simulation efficiency (by removing phenotypes—and hence ODEs—that are negligible) and does not affect the results either.

At the end of each simulation we measure the mean virulence in the free-living parasite population for a given level of specificity, *s*, as:

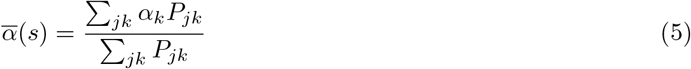

We focus our primary analysis on the effects of the specificity parameter (*s*), the number of susceptibility and infectivity alleles (*n*), and the baseline shedding rate 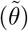 on the mean evolved virulence of the parasite. Default parameter values and ranges (chosen to give representative illustrations of the results) are shown in Table 1. We calculate the percentage change in mean virulence from a baseline of no specificity as:

**Table 1:**
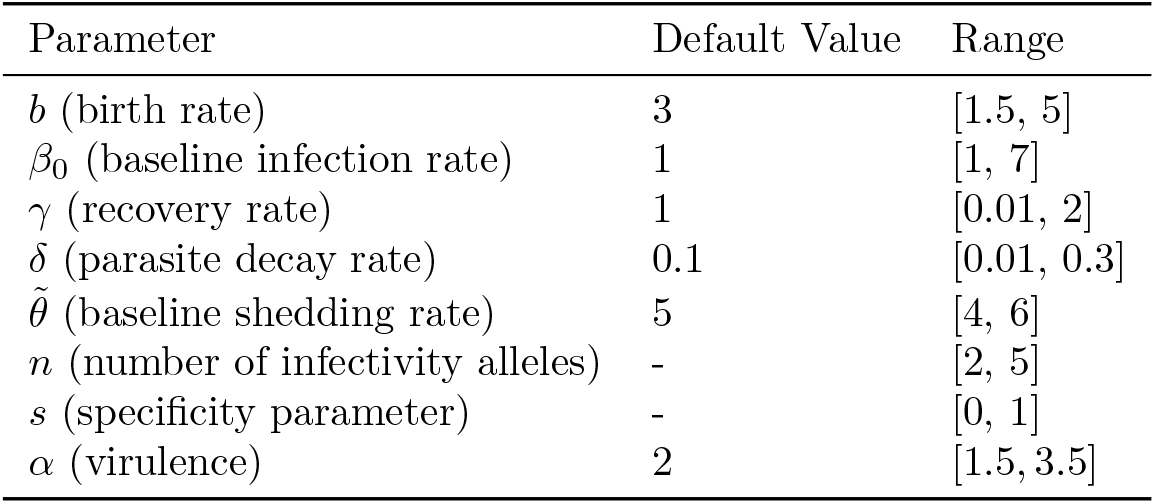
Model parameters, default values, and ranges used in the analysis.

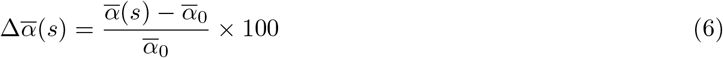

with 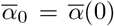. A larger value for 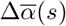 indicates a more substantial impact of specificity on virulence evolution. We also measure the ecological effects of coevolutionary cycling on the mean density of susceptible hosts (assuming a fixed level of virulence) for a given level of specificity as:

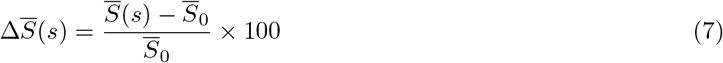

and on mean disease prevalence as:

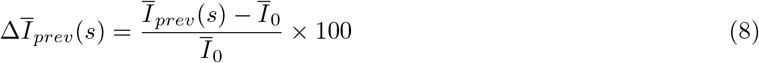

where

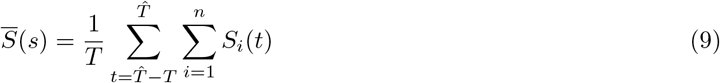

and

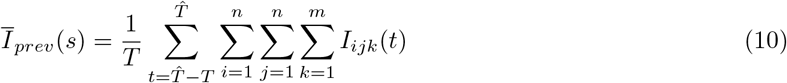

are the mean density of the susceptible host population and mean disease prevalence, respectively, over the time period 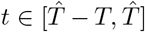, with 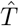 set to a sufficiently large value to avoid transient dynamics and 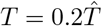 to measure the dynamics over the final 20% of a model run. We define 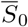 and 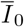 to be the respective values of these measures when there is no specificity (*s* = 0). Thus, 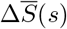 and 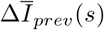 measure how specificity, and hence, coevolutionary cycling affects the mean density of susceptible hosts and mean disease prevalence in the absence of evolution.

## 3 Results

### 3.1 Higher specificity increases coevolutionary cycling and susceptible host density

When there is no specificity (*s* = 0), all entries in *Q*_*ij*_(*n, s*) are identical and therefore all parasites are equally able to infect all hosts, so no coevolutionary cycling occurs (Fig. 1A,D). Introducing specificity (*s >* 0) generates negative frequency-dependent selection, as selection increasingly favours rarer susceptibility alleles in the host population, with parasite infectivity alleles tracking the most common host susceptibility alleles (Fig. 1B,E). Increasing the strength of specificity (*s*) increases the strength of negative frequency-dependent selection, leading to more rapid and higher amplitude cycles in susceptibility and infectivity allele frequencies (Fig. 1C,F). Similarly, increasing the number of susceptibility and infectivity alleles, *n*, tends to produce more rapid and larger amplitude coevolutionary cycling (Figs. 2, S2), with the dynamics eventually becoming chaotic for sufficiently large *n*. Higher levels of specificity–and higher numbers of susceptibility and infectivity alleles–are also associated with an increase in the mean density of susceptible hosts (Fig. 3A) and a decrease in mean disease prevalence (Fig. 3B), with changes in both measures most apparent for intermediate to high levels of specificity.

**Figure 1:**
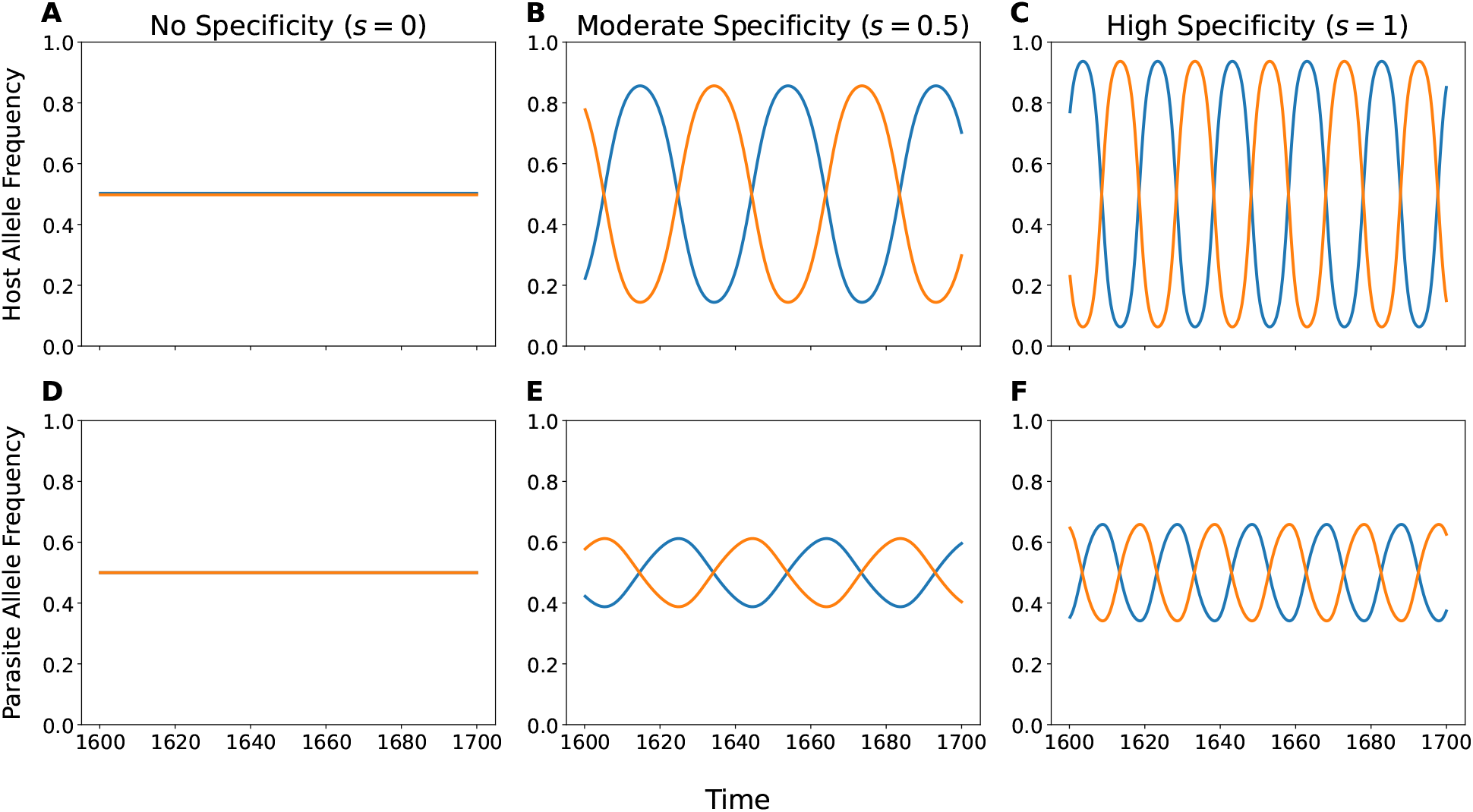
Example susceptibility (top row) and infectivity (bottom row) allele frequency dynamics as specificity, *s*, varies. Colors correspond to different susceptibility or infectivity alleles. All other parameters as described in Table 1, with *n* = 2.

**Figure 2:**
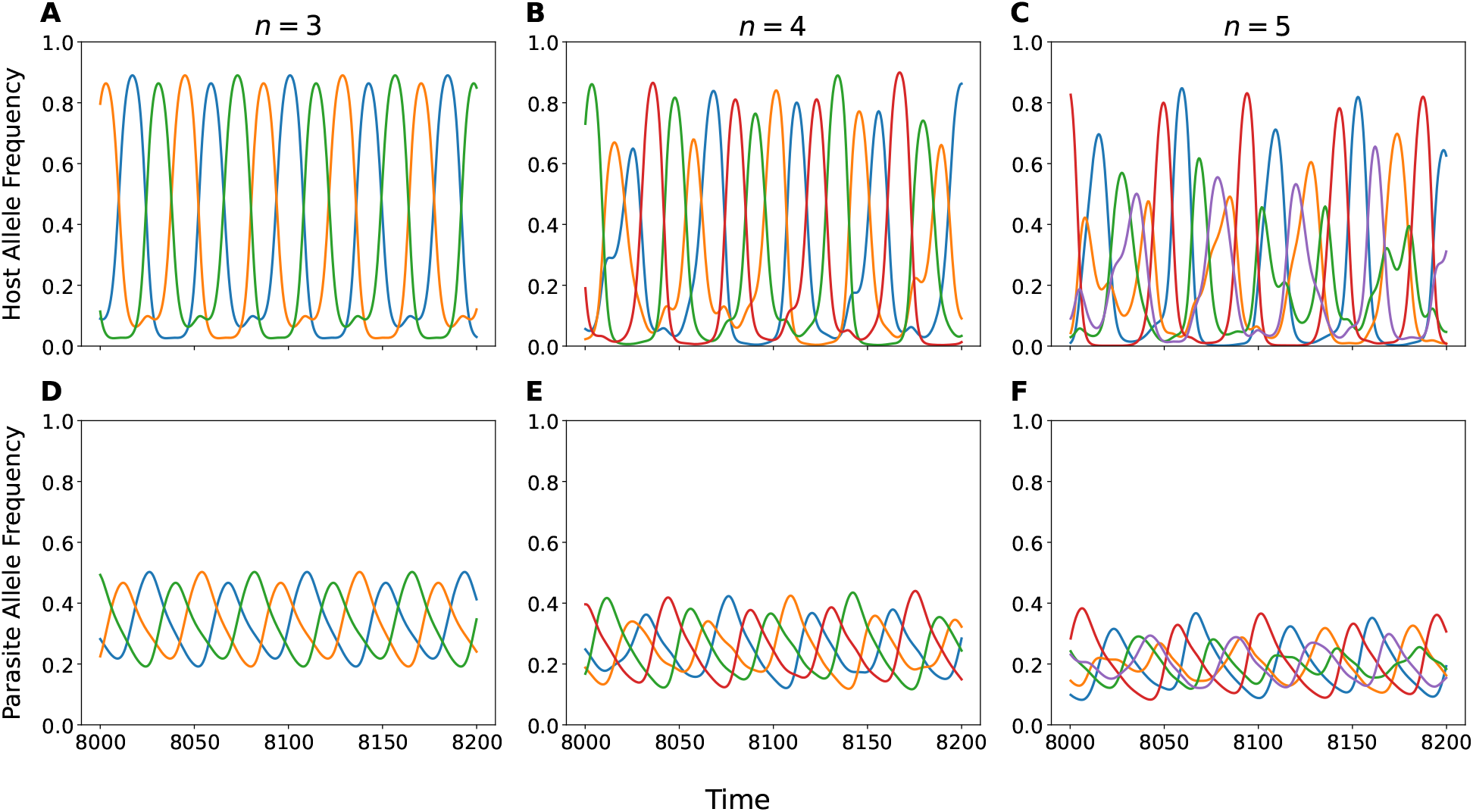
Example susceptibility (top row) and infectivity (bottom row) allele frequency dynamics as the number of alleles, *n*, varies. Colors correspond to different susceptibility or infectivity alleles. All other parameters as described in Table 1, with *s* = 1.

**Figure 3:**
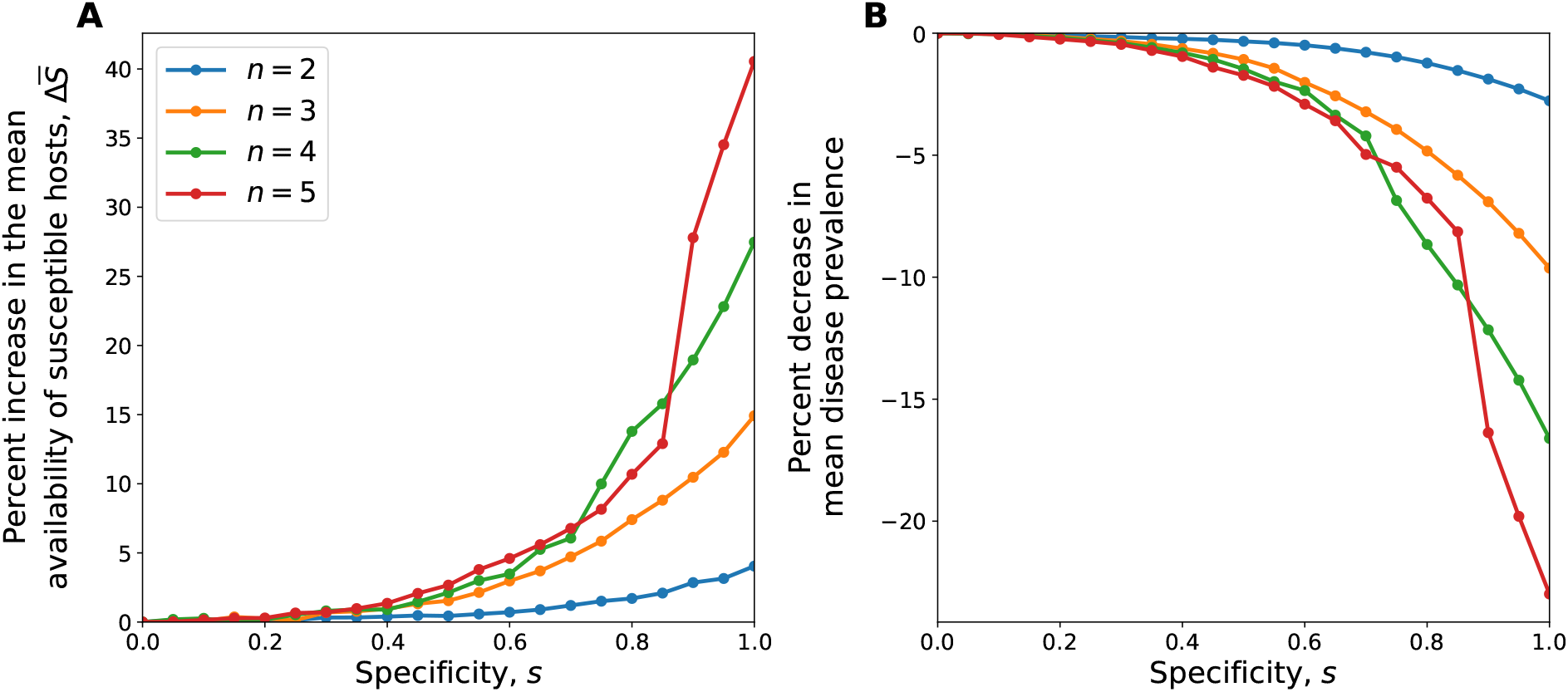
Percent change in the (A) mean density of susceptible hosts, and (B) mean disease prevalence as specificity (*s*) varies for different numbers of susceptibility and infectivity alleles (*n*). Virulence fixed at *α* = 2 and all other parameters as described in Table 1.

### 3.2 More intense coevolutionary cycling selects for higher virulence

As specificity (*s*)—and hence the intensity of coevolutionary cycling—increases, so too does selection for higher virulence (Fig. 4). When specificity is moderate or relatively weak (up to *s ≈* 0.5), coevolutionary cycling has little impact on the evolution of virulence. However, when specificity is relatively strong (*s >* 0.5), there is a significant increase in evolved virulence, with virulence peaking when parasites are only able to infect a single host genotype (*s* = 1). Similarly, as the number of susceptibility and infectivity alleles (*n*) increases, so too do the effects of higher specificity (*s*) on the evolution of virulence (Figs. 4, S3), with greater diversity typically leading to stronger negative frequency-dependent selection for coevolutionary cycling, and in turn, increased selection for virulence (FigS. S2-S3). These results are broadly similar as the baseline shedding rate 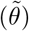 varies, although peak virulence tends to increase with higher baseline shedding rates. Recall that our choice of the baseline infection rate *β*(*n*) and the specificity matrix *Q*_*ij*_(*n, s*) means that the average infection rate at equilibrium does not vary with the strength of specificity (*s*) or the number of infectivity alleles (*n*). This means that effects of these parameters on the evolution of virulence are entirely due to differences in coevolutionary cycles in susceptibility and infectivity allele frequencies.

**Figure 4:**
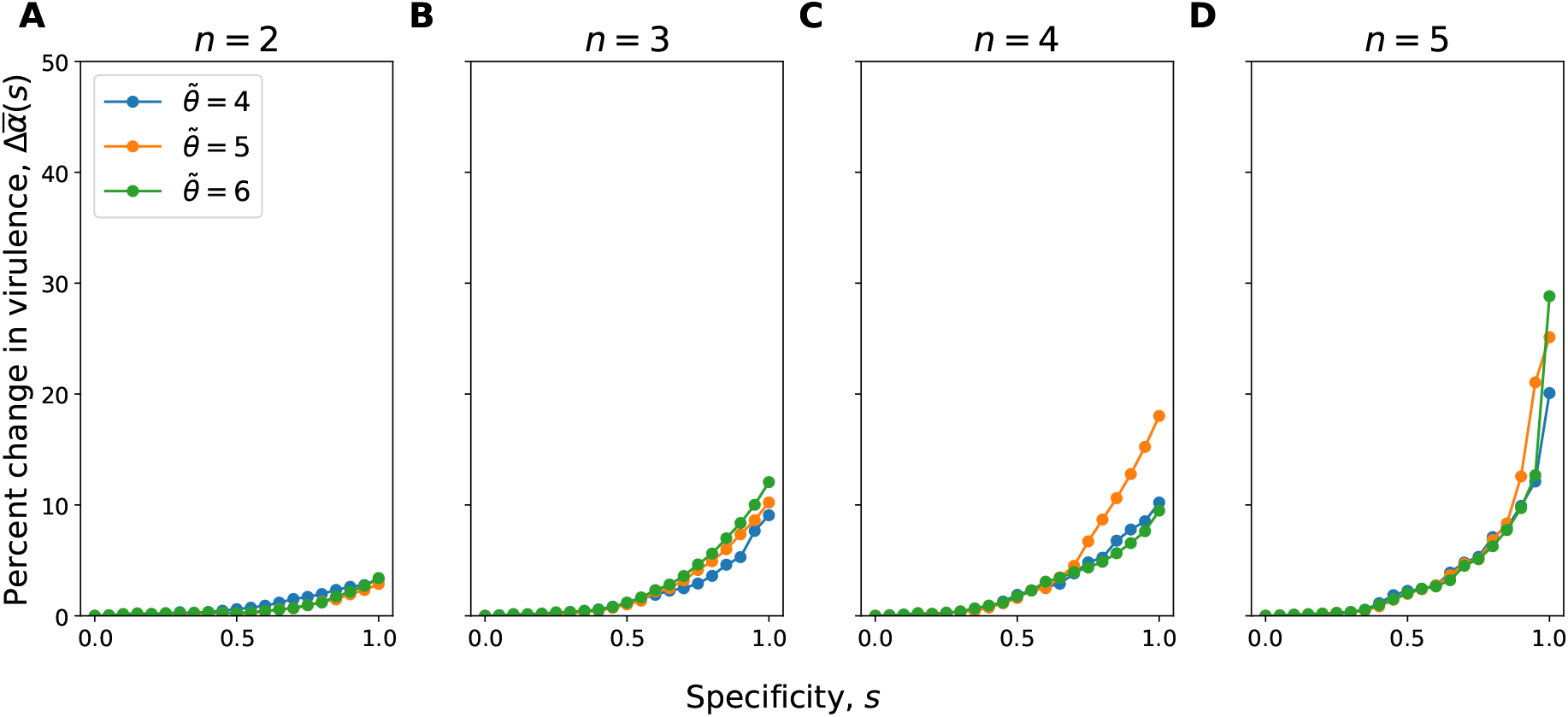
Effects of the specificity parameter (*s*), the number of susceptibility and infectivity alleles (*n*), and the baseline shedding rate 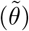 on the evolution of virulence. Percent change in virulence calculated relative to when there is no specificity (*s* = 0). Remaining parameters as specified in Table 1. No variation in the evolved level of virulence was detected between simulations with the same parameters.

Our sensitivity analysis confirmed that these results are robust, with higher specificity always selecting for higher virulence (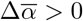 in Fig. 5). Moreover, higher baseline infection rates (*β*_0_), lower recovery rates (*γ*), and higher birth rates (*b*) increase the impact of coevolutionary cycling on the evolution of virulence (i.e., increase the difference in mean evolved virulence from no (*s* = 0) to full (*s* = 1) specificity). In accordance with Lion and Gandon (2022), we find that the change in virulence peaks for intermediate parasite decay rates. In the *Supplementary Material*, we show that our results are also robust to variation in the mutation rate (controlled by *T*_*eco*_, Fig. S4-S5) and when free-living parasites are unable to detach from incompatible hosts (Fig. S6).

**Figure 5:**
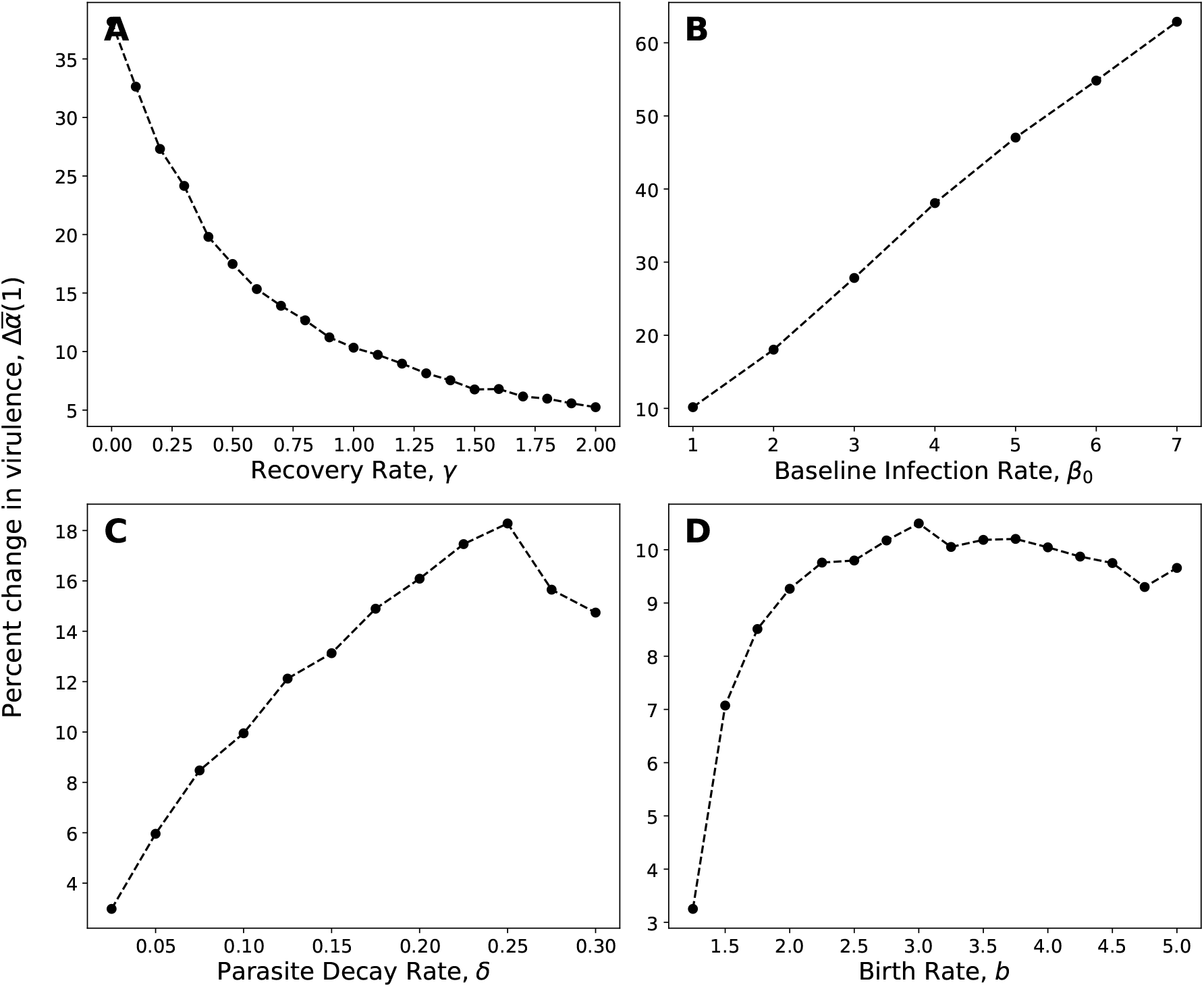
Sensitivity analysis showing the percent change in evolved virulence under full specificity (*s* = 1) relative to no specificity (*s* = 0) with *n* = 3 and 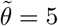. All other parameters as described in Table 1.

## 4 Discussion

Both the evolution of virulence and coevolutionary cycling in allele frequencies have been the focus of intense theoretical and empirical research in recent decades (Buckingham and Ashby 2022). Yet, to the best of our knowledge, this is the first study to explore how coevolutionary cycling—induced by negative frequency-dependent selection—directly affects the evolution of virulence. Using a relatively simple host-parasite model, we have shown that increasing the strength of specificity (*s*) between hosts and parasites generally leads to more intense (higher amplitude and more rapid) coevolutionary cycling in susceptibility and infectivity alleles, which in turn selects for higher levels of virulence. Similarly, increasing the diversity of susceptibility and infectivity alleles (*n*) typically leads to more intense coevolutionary cycling and the evolution of greater virulence. Our findings are robust to variation in all model parameters and when parasites are unable to detach from incompatible hosts, suggesting this is likely to be a fairly general phenomenon.

The specificity matrix, *Q*_*ij*_(*n, s*), and transmission rate, *β*(*n*), are key features of our model, which we constructed in such a way as to prevent the strength of specificity (*s*) or the number of infectivity alleles (*n*) affecting the mean infection rate for parasites at equilibrium. This means that any differences in evolved virulence can be fully attributed to patterns of coevolutionary cycling in susceptibility and infectivity alleles. While neither specificity nor the diversity of the populations affect the ecological equilibrium in our model, they do affect both the mean density of susceptible hosts and mean disease prevalence when the populations cycle (given a fixed level of virulence). Specifically, the mean density of susceptible hosts generally increases with *s* and *n*, while mean disease prevalence falls. These ecological patterns are consistent with the dilution effect, where greater host diversity tends to reduce disease prevalence (Keesing and Ostfeld 2021). However, the dilution effect also predicts that parasites will evolve lower virulence due to reduced transmission opportunities (White et al. 2020). This does not occur in our model because we balance the effects of increased host diversity on parasite transmission by setting the transmission rate to be proportional to the number of host genotypes. Instead, we typically see selection for higher virulence as host diversity increases due to coevolutionary cycling.

Greater specificity and diversity in our model lead to more extreme cycles in allele frequencies, with each host allele reaching higher peak abundances, which increases the density of susceptible hosts on average. We believe that more intense coevolutionary cycling selects for higher virulence in our model because this cycling increases the mean density of susceptible hosts. This in turn allows for higher exploitation rates by parasites as susceptible hosts are more abundant. In contrast, cycling induced by seasonal forcing has previously been shown to have no effect on the evolution of virulence (unless virulence is density-dependent), as there is no impact on the mean density of susceptible hosts (Donnelly et al. 2013). We therefore strongly suspect that the reason more intense coevolutionary cycling selects for higher virulence is due to its effects on the mean density of susceptible hosts.

Our results also have implications for understanding the evolution of virulence in specialists and generalists. It has been argued that generalist parasites could either be more or less virulent than specialists depending on the context (Leggett et al. 2013), with empirical studies finding both outcomes are possible (Garamszegi 2006; Agudelo-Romero and Elena 2008; Torres-Barceló et al. 2024). Our study suggests that specialists may indeed evolve to be more virulent than generalists, but crucially, this occurs in our model due to population dynamics (variation in the density of susceptible hosts) as opposed to specialists being better adapted than generalists to exploit their hosts.

The model studied herein assumes a free-living parasite population. Many parasites exhibit free-living stages (e.g. bacteriophages, protozoa such as Giardia, and helminths), but are less commonly studied in theoretical evolutionary biology (although see for example, Lion and Gandon (2022)). For the purposes of our study, which sought to demonstrate how coevolutionary cycling *per se* can drive virulence evolution, a free-living parasite stage has the advantage of being able to sustain coevolutionary cycling indefinitely. Comparable models without a free-living parasite stage typically produce only damped oscillations and so these transient dynamics will not have a long-term impact on virulence evolution. However, in the limit of rapid free-living parasite dynamics, one can make a quasi-steady-state approximation to separate fast parasite and slow host dynamics, so that the free-living parasite model converges to a model without free-living parasites. Furthermore, models of infectious diseases can exhibit sustained oscillations due to additional factors such as stochasticity (Kuske et al. 2007) or seasonality (Donnelly et al. 2013), and so we would also expect to see similar long-term effects of coevolutionary cycling on the evolution of virulence in real populations even in the absence of a free-living parasite stage.

Our study is closely related to work by Kirchner and Roy (2002) and Gandon et al. (2002), but differs in key aspects from both. Kirchner and Roy (2002) explored the effects of specificity and coevolutionary cycling on pathogen coexistence, but their model was also limited to two host and two parasite genotypes, and virulence was not allowed to freely evolve. Gandon et al. (2002) showed that lower virulence can be selected for in a model of coevolution, but the populations were assumed to be at equilibrium. In contrast, our study explicitly focuses on coevolution as a dynamic, non-equilibrium process, with persistent fluctuations in allele frequencies driven by negative frequency-dependent selection. Furthermore, Gandon et al. (2002) considered a different framework of infection genetics known as the Gene-for-Gene (GFG) model, which assumes there is variation among parasites (hosts) in the range of hosts (parasites) that can be infected (resisted) (Flor 1956; Frank 1993). Here, we assumed that parasites (hosts) only differ in which hosts (parasites) that they specialise on infecting (resisting). This is a generalisation of the Matching Alleles (MA) framework of infection genetics, which is based on self/non-self recognition mechanisms (Frank 1993; Grosberg and Hart 2000) and is considered to be at the opposite end of a spectrum with the GFG model (Agrawal and Lively 2002) (or, in multilocus systems, a subset of the GFG model (Ashby and Boots 2017)). We focused on MA-like models to ensure that variations in specificity did not affect the average infection rate of parasites, and hence any changes in the evolution of virulence could be fully attributed to the effects of specificity (and hence, coevolutionary cycling). Future work should consider how coevolutionary cycling in asymmetrical infection genetics frameworks such as GFG-like models affects the evolution of virulence.

A growing body of evolutionary theory shows that non-equilibrium dynamics can profoundly affect parasite evolution (Best and Ashby 2023). In particular, several studies have shown that fluctuations in population densities induced by forces other than coevolution have can influence the evolution of virulence. For example, Donnelly et al. (2013) showed that seasonally forced host reproduction leads to the evolution of higher density-dependent virulence, and Hite and Cressler (2018) found that emergent—as opposed to seasonally forced—fluctuations in host reproduction can lead to the evolution of extreme levels of virulence (high or low) due to bistability. Both of these studies assumed a virulence-transmission trade-off similar to the one in our model, but MacDonald et al. (2022) showed that even in the absence of such a trade-off, fluctuations in host availability can affect the evolution of virulence. Finally, Lion and Gandon (2022) applied a novel method for class-structured populations to analyse the effects of fluctuating environments on parasite evolution, showing that fluctuating environments can have contrasting effects depending on the context: fluctuating environments select for more virulent long-lived parasites, but reduce selection for virulence when imperfect vaccines are used. Together, these studies highlight that we cannot necessarily extrapolate predictions for parasite evolution from constant to fluctuating environments.

Detecting coevolutionary cycling empirically is challenging, and is often achieved through time-shift experiments (Gandon et al. 2002; Gaba and Ebert 2009). Some of the strongest evidence for coevolutionary cycling comes from experimental evolution of bacteria and phages (Gómez and Buckling 2011), observations of water snails and trematodes (Dybdahl and Lively 1998), and planktonic crustaceans and bacteria (Decaestecker et al. 2007). As far as we are aware, the effects of coevolutionary cycling *per se* on the evolution of parasite virulence has yet to be explored in these systems. Experimentally manipulating the specificity of the parasite is likely difficult, but our findings suggest that varying the number of host and parasite genotypes (i.e., diversity of susceptibility and infectivity alleles, *n*) should lead to selection for different levels of virulence (provided the overall transmission rate does not vary), with greater diversity selecting for higher virulence.

The effects of coevolution on other life-history traits has received surprisingly little attention given the large body of theoretical literature on coevolution (Buckingham and Ashby 2022). Our results show that coevolutionary cycling in allele frequencies can indeed have profound effects on the evolution of virulence, with specialist parasites undergoing more intense coevolutionary cycling and hence evolving to be more virulent than generalists.

