## Supplementary Material for "Coevolutionary cycling in allele frequencies and the evolution of virulence"

### S1 Simulations

#### S1.1 Overview

To analyze the impacts of coevolutionary cycles on the evolutionary dynamics of virulence, we run hybrid (deterministic and stochastic) simulations in C++ (source code available at `https://github.com/dxanielkm/Ashby_and_Kim_2025/tree/main`). Each simulation is initialized with all susceptibility and infectivity alleles present at arbitrary population densities, and a single virulence phenotype for all parasites. The trait space for virulence is discretized into $m = 21$ equally spaced traits, with boundaries as specified in Table 1 in the main text. The boundaries are chosen based on preliminary simulations to ensure that the parasite does not reach either boundary when virulence evolves. We initialize all parasites at the lower bound of this trait space.

The simulations are separated into ecological and evolutionary timescales. Within each evolutionary time step, the ecological dynamics of extant hosts and parasites are described by equations (2)-(4) in the main text. The ecological dynamics are progressed deterministically for a fixed number of time units, $T_{eco}$. The duration of ecological dynamics effectively controls the mutation rate for the virulence locus, with smaller values of $T_{eco}$ corresponding to a higher mutation rate (we set $T_{eco} = 400$ throughout). Any population class that is below an extinction threshold of $\epsilon = 10^{-9}$ at the end of the ecological time period is set to 0. The duration for the ecological dynamics and the extinction thresholds are fairly arbitrary, with preliminary simulations confirming that a wide range of values produce similar results, provided $T_{eco}$ is not too small and $\epsilon$ is not too large. The ecological dynamics are then paused and we introduce a rare mutant parasite, $\alpha_{mut}$, with a slightly different virulence phenotype to one of the resident phenotypes, $\alpha_{res}$. The progenitor for the mutant is chosen randomly based on a weighted probability of parasite frequencies. The ecological dynamics with the new mutant are then progressed again by $T_{eco}$ time units and the process repeats for a total of $T_{evo} = 4,000$ evolutionary time steps (this duration was sufficient for convergence to an evolutionarily stable level of virulence in all simulations).

Example simulation trajectories are shown in Fig. S1 for three different levels of specificity ($s \in \{0, 0.5, 1\}$) and three ecological time periods ($T_{eco} \in \{200, 400, 800\}$). The value of $T_{eco}$ only affects the effective mutation rate and therefore only affects the time taken to reach the evolutionarily stable level of virulence, rather than the level of virulence that is evolutionarily stable.

#### S1.2 Mutation algorithm

We use the following algorithm to introduce a rare mutant parasite at each evolutionary time step:

1. Calculate the cumulative distribution function (CDF), $C_v$, of length $mn$ with $v$th entry given by:

$$C_v = \sum_{u=1}^{v} P_{j_u k_u} \tag{S1}$$

where $u$ indexes over all combinations of infectivity ($j_u \in \{1, 2, ..., n\}$) and virulence ($k_u \in \{1, 2, ..., m\}$) alleles.

2. Select the progenitor for the mutation by generating a random number, $r$, uniformly distributed between 0 and $C_{mn}$, and finding the smallest $q$ such that $C_q \geq r$. This ensures that genotypes with a higher frequency in the population are more likely to be the source of the next virulence mutation.

3. The mutant virulence trait $\alpha_{k_{mut}}$ is chosen adjacent (subject to boundaries) to the parent trait $\alpha_{k_q}$ such that:

$$k_{mut} = \begin{cases} \max(1, k_q - 1), & \text{with probability } 0.5 \\ \min(m, k_q + 1), & \text{with probability } 0.5 \end{cases}$$

   The corresponding infectivity allele ($j_q$) remains unchanged.

4. Transfer a small fraction $\eta \ll 1$ of the progenitor population, $P_{j_q k_q}$ to the mutant population, $P_{j_q k_{mut}}$, such that:

$$P_{j_q k_{mut}} \leftarrow P_{j_q k_{mut}} + \eta P_{j_q k_q}, \quad P_{j_q k_q} \leftarrow P_{j_q k_q}(1 - \eta) \tag{S2}$$

#### S2 Effects of the number of susceptibility and infectivity alleles, $n$, on eco-evolutionary dynamics

In the main text, we show that increasing the number of susceptibility and infectivity alleles, $n$ leads to higher evolved levels of virulence. To further explore the effects of host and parasite diversity on model dynamics, here we analyze how the amplitude and frequency of coevolutionary cycles varies with larger $n$, and in turn, how virulence evolves.

For each host allele, we analyzed the maximum amplitude of coevolutionary cycles in allele frequencies by first locating the local maxima and minima with a minimum prominence of $10^{-3}$ using a standard peak-finding algorithm in Python. We then pruned extrema separated by less than $\Delta t_{\min} = 0.1$ to avoid spurious results. Finally, we computed the amplitudes of coevolutionary cycles as the difference between consecutive extrema and recorded the maximum over all amplitudes computed. Similarly, to calculate the maximum frequency of coevolutionary cycles, we identified the peak times for each host allele and applied the same prominence and pruning methods described above for the amplitude calculation. We then pooled peak times across the $n$ host alleles and computed the frequencies as $1/\Delta t$ (where $\Delta t$ is the difference in peak times).

The results of this analysis are shown in Fig. S2, with both the maximum amplitude and frequency of coevolutionary cycles increasingly monotonically with $n$. In turn, we see virulence generally evolving to higher values as $n$ increases (Fig. S3), but with $n$ having diminishing effects for larger numbers of alleles.

#### S3 Mutation rate

The mutation rate is controlled by the duration of ecological dynamics, $T_{eco}$, with shorter ecological windows corresponding to faster mutation rates. Crucially, the mutation rate does not affect the long-term evolution of virulence, only the rate of adaptation and the breadth of the distribution of phenotypes around mean virulence. In the main text we fix $T_{eco} = 400$, which was chosen to provide a reasonable mutation supply while maintaining computational efficiency. Here, we demonstrate that increasing or decreasing the mutation rate by shortening ($T_{eco} = 200$; Fig. S4) or lengthening ($T_{eco} = 800$; Fig. S5) the duration of ecological dynamics between mutations has no qualitative impact on the results.

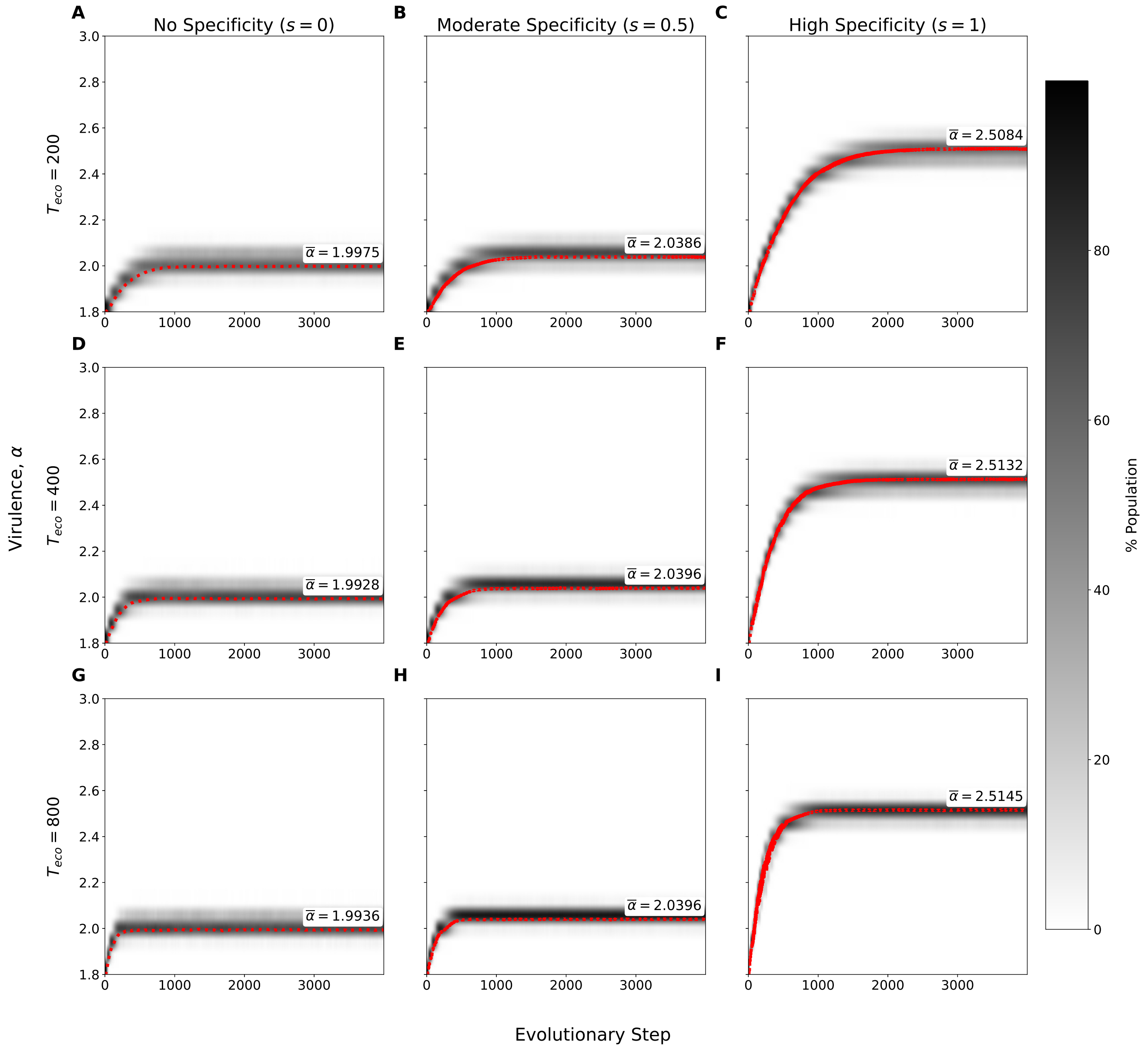


Figure S1: Example evolutionary trajectories showing average virulence over all infectivity alleles, under varying levels of specificity ($s \in \{0, 0.5, 1\}$) and varying levels of ecological time ($T_{eco} \in \{200, 400, 800\}$). Shaded regions show distribution of virulence phenotypes averaged over all infectivity alleles, with darker regions indicating more common virulence phenotypes. Red lines indicate the mean level of virulence in the parasite population. Remaining parameters specified in Table 1 of the main text.

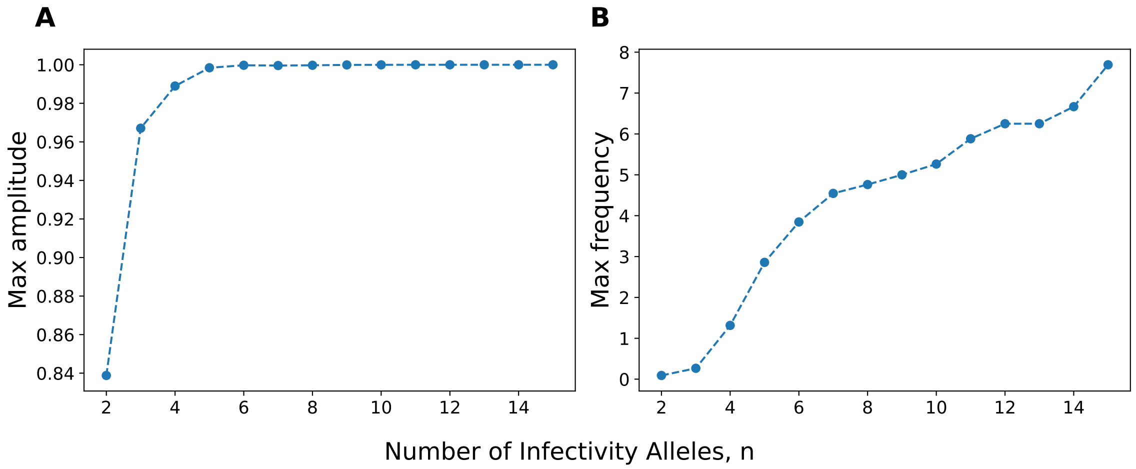


Figure S2: Effects of the number of infectivity alleles ($n$) on coevolutionary cycling. Specificity parameter was set to $s = 1$ for all model runs. Remaining parameters specified in Table 1 of the main text.

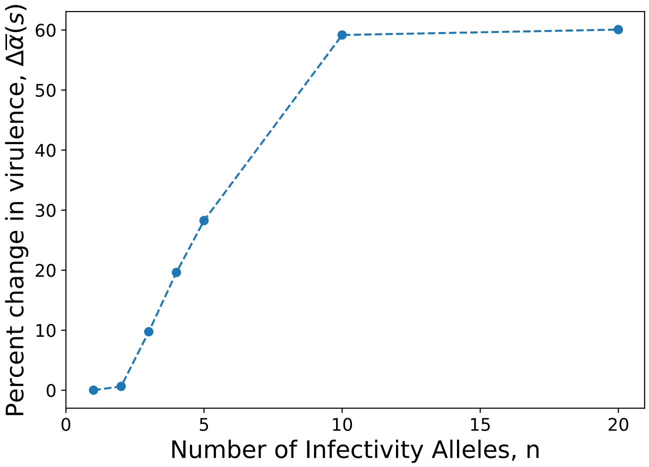


Figure S3: Effects of the number of susceptibility and infectivity alleles ($n$) on the evolution of virulence. Percent change in virulence calculated relative to when there is no specificity ($s = 0$). Remaining parameters specified in Table 1 of the main text.

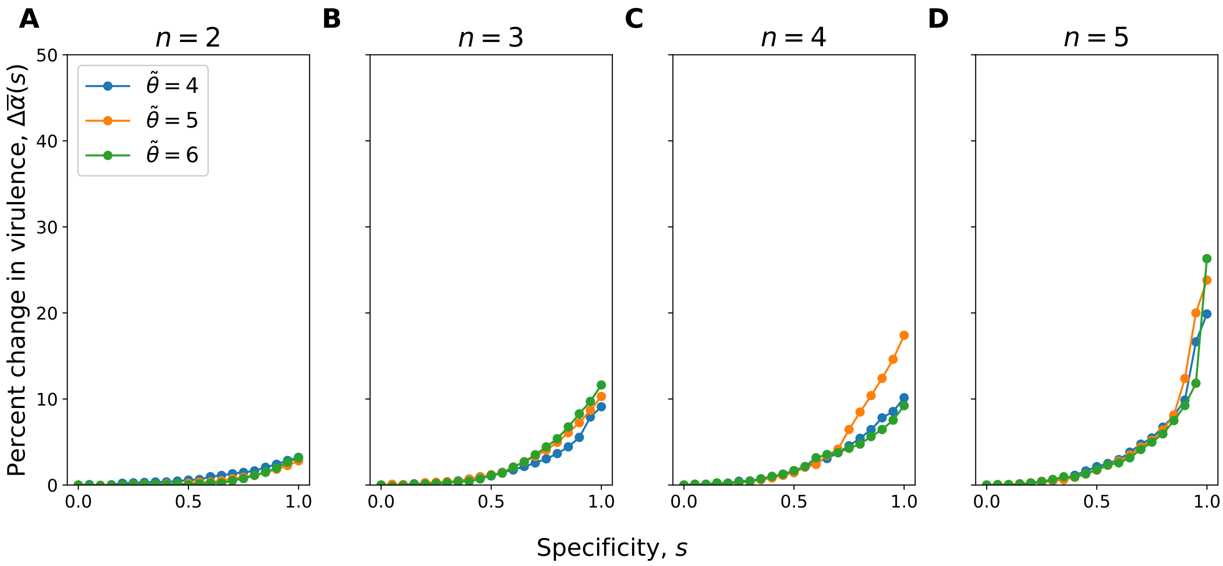


Figure S4: Effects of the specificity parameter ($s$), the number of susceptibility and infectivity alleles ($n$), and the baseline shedding rate ($\tilde{\theta}$) on coevolutionary cycling with all parameters as in Fig. 4 of main text, except $T_{eco} = 200$.

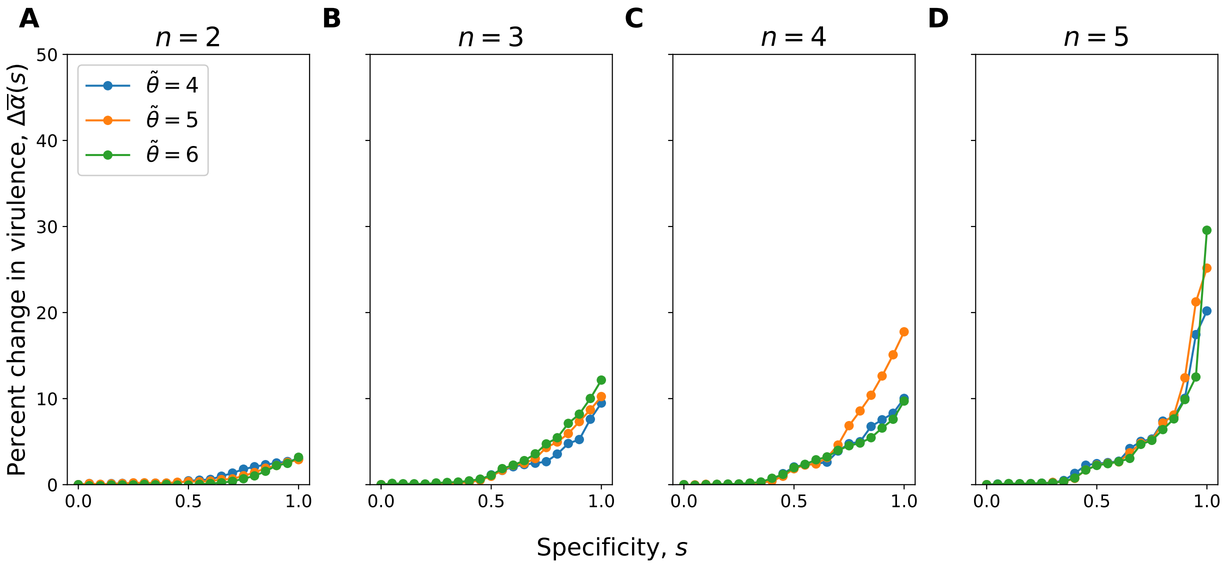


Figure S5: Effects of the specificity parameter ($s$), the number of susceptibility and infectivity alleles ($n$), and the baseline shedding rate ($\tilde{\theta}$) on coevolutionary cycling with all parameters as in Fig. 4 of main text, except $T_{eco} = 800$.

#### S4 Irreversible Detachment Model

In the main text we assume that free-living parasites that attempt but fail to infect hosts due to incompatibility do not suffer any negative consequences (i.e., they can detach from the incompatible host and potentially infect a compatible host in future without penalty). Here, we explore a variation of the model where free-living parasites that attempt to infect incompatible hosts are killed. Hence, any attempted attachment removes the parasite from the environment, regardless of whether infection was successful or not. We refer to this as the "Irreversible Detachment Model". The only term that requires modification is the final term in equation (4) from the main text, where we now remove the specificity matrix $Q_{ij}(n,s)$:

$$\frac{dP_{jk}}{dt} = \theta_k \sum_{i=1}^{n} I_{ijk} - \delta P_{jk} - \sum_{i=1}^{n} \beta(n) P_{jk} S_i \tag{S3}$$

As shown in Fig. S6, the impact of specificity and the number of infectivity alleles on the evolution of virulence is qualitatively similar to the original model (Fig. 4).

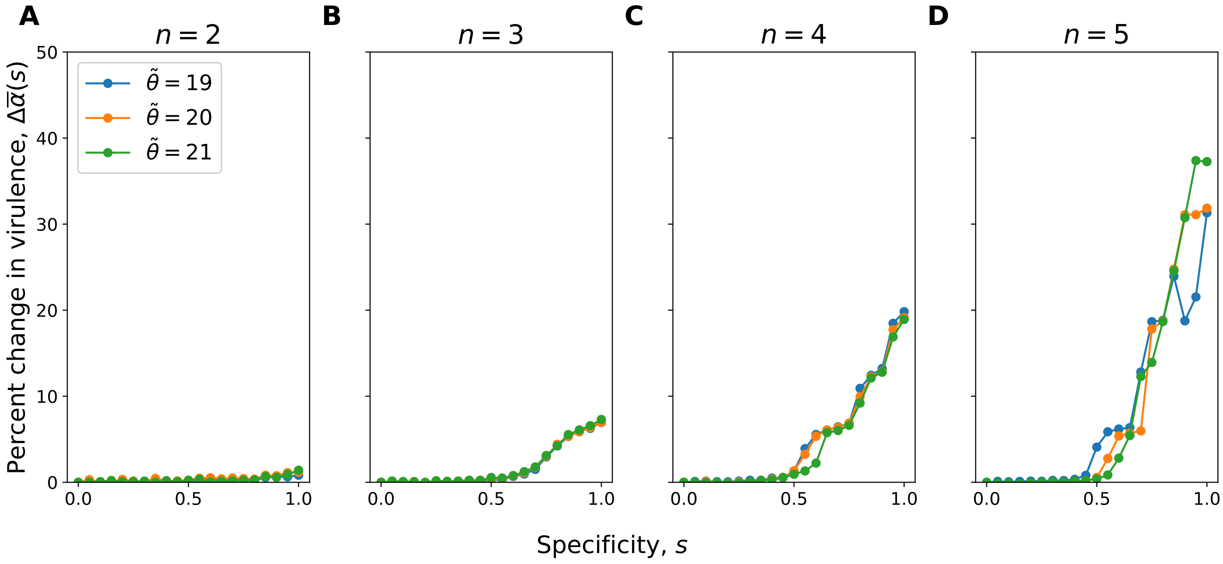


Figure S6: Effects of the specificity parameter ($s$), the number of susceptibility and infectivity alleles ($n$), and the baseline shedding rate ($\tilde{\theta}$) on the evolution of virulence in the Irreversible Detachment Model. Percent change in virulence calculated relative to when there is no specificity ($s = 0$). Remaining parameters as specified in Table 1 of the main text.
